# Maturation of complex synaptic connections of layer 5 cortical axons in the posterior thalamic nucleus requires SNAP25

**DOI:** 10.1101/2020.05.20.093070

**Authors:** Shuichi Hayashi, Anna Hoerder-Suabedissen, Emi Kiyokage, Catherine Maclachlan, Kazunori Toida, Graham Knott, Zoltán Molnár

## Abstract

Synapses are able to form in the absence of neuronal activity, but how is their subsequent maturation affected in the absence of regulated vesicular release? We explored this question using 3D electron microscopy and immuno electron microscopy analyses in the large, complex synapses formed between cortical sensory efferent axons and dendrites in the posterior thalamic nucleus. Using a *Snap25* conditional knockout we found that during the first two postnatal weeks the axonal boutons emerge and increase in the size similar to the control animals. However, by P18, when an adult-like architecture should normally be established, axons were significantly smaller with 3D reconstructions showing that each *Snap25*-cko bouton only forms a single synapse with the connecting dendritic shaft. No excrescences from the dendrites were formed, and none of the normally large glomerular axon endings were seen. These results show that activity mediated through regulated vesicular release from the presynaptic terminal is not necessary for the formation of synapses, but it is required for the maturation of the specialised synaptic structures between layer 5 corticothalamic projections in Po.

## Introduction

Synaptic development involves coordinated morphological changes of presynaptic and postsynaptic neurites such as formation of boutons and spines. Presynaptic mechanisms for those processes include neuronal activity and molecular signals that target postsynaptic dendrites (Andreae and Burrone, 2018; Bleckert and Wong, 2011; Favuzzi and Rico, 2018; Shen and Cowan, 2010). The role of neuronal activity in synapse formation in the brain has been extensively studied with different approaches. Blocking synaptic transmission by knocking out the gene for mammalian uncoordinated (Munc)-13, Munc-18 or the SNARE-complex protein Synaptosome Associated Protein 25 (SNAP25) does not affect synapse formation in embryonic stages (Augustin et al., 1999; Varoqueaux et al., 2002; Verhage et al., 2000; Washbourne et al., 2002). More recent studies have supported the idea that synaptic transmission is dispensable for the formation and maintenance of synapses or spines in postnatal hippocampal CA1 by analysing neurons with neonatal deletion of subunits of the NMDAR and AMPA receptors (Lu et al., 2013), organotypic slice cultures of double knockout of *Munc13-1* and *13-2* (Sigler et al., 2017) or Emx1-driven tetanus toxin (TeNT) expressing brains (Sando et al., 2017). However, this only addresses the effect of silencing cortical neurons in the cortex and hippocamus. How does it affect synapse morphogenesis and maturation in regions that receive large and powerful cortical inputs such as the thalamic glomerular type connections in the thalamus (Groh et al., 2008; Hoerder-Suabedissen et al., 2018; Hoogland et al., 1991; Maclachlan et al., 2018) ?

Corticofugal projections from cortical layer 5 provide major descending projections to subcortical areas, and those from the motor, sensory and visual cortices have branches into ‘higher-order (associate)’ thalamic nuclei (Deschenes et al., 1994; Kita and Kita, 2012). It has been proposed that those layer 5 corticothalamic projections mediate trans-thalamic communication of different cortical areas (Saalmann, 2014; Sherman and Guillery, 2011). Layer 5 corticothalamic projections from somatosensory cortex in mice are known to have characteristic large boutons (giant boutons) in the posterior thalamic nucleus (Po), one of the higher-order thalamic nuclei (Groh et al., 2008; Hoerder-Suabedissen et al., 2018; Hoogland et al., 1991; Maclachlan et al., 2018). Unlike layer 6 corticothalamic projections, which have always small boutons, layer 5 corticothalamic projections from primary somatosensory (S1) cortex to Po provide powerful inputs to postsynaptic neurons mediated through ionotropic glutamate receptors (Groh et al., 2008; Reichova and Sherman, 2004). This morphological and electrophysiological relationship suggests that the large bouton structure of layer 5 is essential for faithfully transferring powerful inputs to thalamic neurons (Sherman and Guillery, 2011).

*Snap25* null mice show normal brain development during embryonic stages, but the neonatal lethality hampers the study on the role of SNAP25 in synaptic development during postnatal periods (Molnár et al., 2002; Washbourne et al., 2002). Our recent study using a cortical layer 5 Cre driver, *Rbp4-Cre*, and *Snap25* conditional knockout (*Snap25* cKO) mice has demonstrated that *Snap25* deletion in cortical layer 5 does not affect their axon projections and targeting during development but causes degeneration of their axons in the adult brain (Hoerder-Suabedissen et al., 2019). The utilisation of *Rbp4-Cre*-driven *Snap25* cKO has some advantages to study the development of synapses in the corticothalamic system. First, *Rbp4-Cre* starts to be expressed at late embryonic stages (Grant et al., 2016), which is early enough to understand how layer 5 specialised giant boutons develop during the postnatal periods. Second, since *Snap25* removal does not affect the ingrowth of layer 5 corticothalamic projections into thalamic nuclei (Hoerder-Suabedissen et al., 2019), we can specifically address the effect of *Snap25* ablation on the specialised synaptic development. Therefore, using *Rbp4-Cre* driven ablation of *Snap25* in this study, we found that layer 5 giant boutons, and their synapses in Po, develop during the second and third postnatal weeks. However, although presynaptic SNAP25 is dispensable for the initial formation of these synapses, their subsequent maturation which includes the growth of boutons and protrusion of multiple excrescences from thalamic dendrites appears to be impeded. Our results, therefore, suggest that the presynaptic control of regulated vesicular release by SNAP25 is essential for establishing the specialised synaptic connections of layer 5 corticothalamic projections.

## Results

### Layer 5 giant boutons in posterior thalamic nucleus develop in the second and third postnatal weeks

Previous studies using injection of anterograde tracers, or Thy1 promotor-driven fluorescence labelling, have shown that layer 5 neurons from S1 cortex have giant boutons in Po of adult brains (Groh et al., 2008; Hoerder-Suabedissen et al., 2018; Hoogland et al., 1991). To study the synaptic development of layer 5 corticothalamic projections, we used the *Rbp4-Cre;tdTomato* mouse line, whose tdTomato expression starts in a subset of layer 5 neurons at late embryonic stages (Grant et al., 2016; Hoerder-Suabedissen et al., 2019). This allows us to visualise their corticothalamic projections innervating Po in the early postnatal period. We first examined boutons that were positive for *Rbp4-Cre*-driven tdTomato fluorescence (tdTom+) in Po of *Rbp4-Cre*;*Ai14* adult brains. Although *Rbp4-Cre* is widely expressed in cortical layer 5, the somatosensory cortex including S1 is the major source of projections into Po (Allen Brain Connectivity Atlas). Consistent with this, injections of Cre-dependent AAV-eGFP virus to S1 cortex showed that eGFP-positive (eGFP+) axons project to Po (Fig. 1A) and they had large boutons, which colocalised with the presynaptic marker vesicular glutamate transporter 1 (VGluT1), in Po (Fig. 1B). Pre-embedding electron microscopy showed that eGFP+ boutons in Po contain multiple excrescences from Po dendrites (Fig. 1C, D). The cross-sectional area of boutons in randomly selected sections containing Po in adult brains was 2.0±1.0 μm^2^ (mean±s.d., Fig. 1E, n=26 boutons from three brains). The ultrastructural profile and the size of the boutons are comparable to those of *Rbp4-Cre*+ large boutons in Po in our previous study (Hoerder-Suabedissen et al., 2018). We previously reported that *Rbp4-Cre;Ai14* axons also form small boutons in Po and other thalamic nuclei (Hoerder-Suabedissen et al., 2018), but they were not subject of these investigations which specifically targeted the large boutons in Po with specialised synapses that could be easily identified with Cre-dependent viral tracing (AAV-eGFP) or with tdTomato (Figure 1A and B). These results suggest that the presynaptic bouton structure containing multiple dendritic excrescences are a general profile of *Rbp4-Cre* positive large boutons in Po including those specifically derived from S1 cortex.

**Figure 1.**
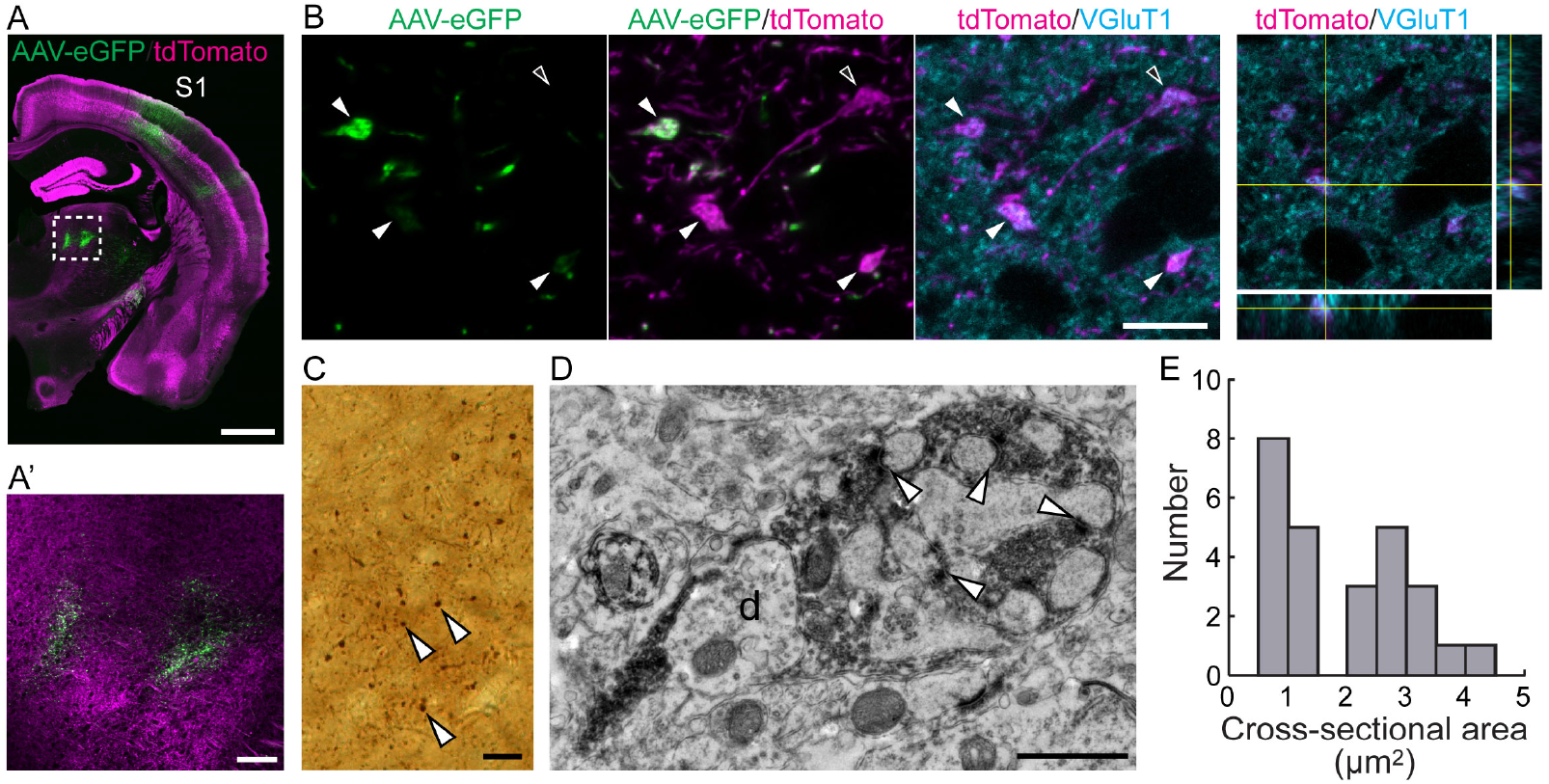
*Rbp4-Cre*+ layer 5 corticothalamic projections from S1 have giant boutons in adult Po. **A**, **B**, Laser-scanning confocal microscopy images of an *Rbp4-Cre;Ai14* adult mouse brain injected with AAV-eGFP in S1 cortex. **A**, Tiled image of *Rbp4-Cre*-dependent expression of AAV-eGFP injected into S1 cortex. **A’**, Higher magnification of the boxed region in **A**, which indicates *Rbp4-Cre* driven eGFP+ (green) and tdTom+ (magenta) axon terminals in Po. **B**, Typical examples of AAV-eGFP+ and/or tdTom+ boutons in Po, which are colocalised with the presynaptic marker VGluT1. The left three images show maximum intensity projections of part of a z-stack (1.0 μm out of 6 μm in total thickness) for each colour shown in top. Orthogonal views at the point of one of the boutons shown in the right. Filled arrowheads and an open arrowhead indicate AAV-eGFP+;tdTom+ boutons and an AAV-eGFP-negative;tdTom+ bouton, respectively. **C**, Light microscopy image of DAB-stained eGFP+ boutons in Po. Arrowheads indicate examples of eGFP+ boutons. **D**, Pre-embedding immuno-electron microscopy against eGFP, with image taken in Po thalamus. White arrowheads point to synapses, ‘d’ denotes dendrite. **E**, Distribution histogram of the cross-sectional area of eGFP+ boutons in Po measured in electron micrographs. The presented net bouton area excludes the dendritic excrescences. N = 26 boutons from three brains were analysed. Scar bars, 1 mm in **A**, 100 μm in A’, 10 μm in **B**, 20 μm in **C**, 1 μm in **D**.

We next examined the development of synapses of tdTom+ axons in Po. tdTom+ axons are shown to innervate Po in the first postnatal week (Grant et al., 2016; Hoerder-Suabedissen et al., 2019). tdTom+ boutons, which colocalised with VGluT1, were detected in Po at P8. To measure the size of boutons in fluorescent microscopy images, z-stack images covering the boutons of interest were projected onto a single plane with maximum intensity projection, and the area of the projected image (projected area) was analysed. The size of the tdTom+ boutons increased by P21 (Fig. 2A,B). The size of the tdTom+ boutons at P21 and P28 was the same as that of the adult (Fig. 2B). These results indicate that the size of *Rbp4-Cre* positive layer 5 boutons in Po increases during the second and third postnatal weeks.

**Figure 2.**
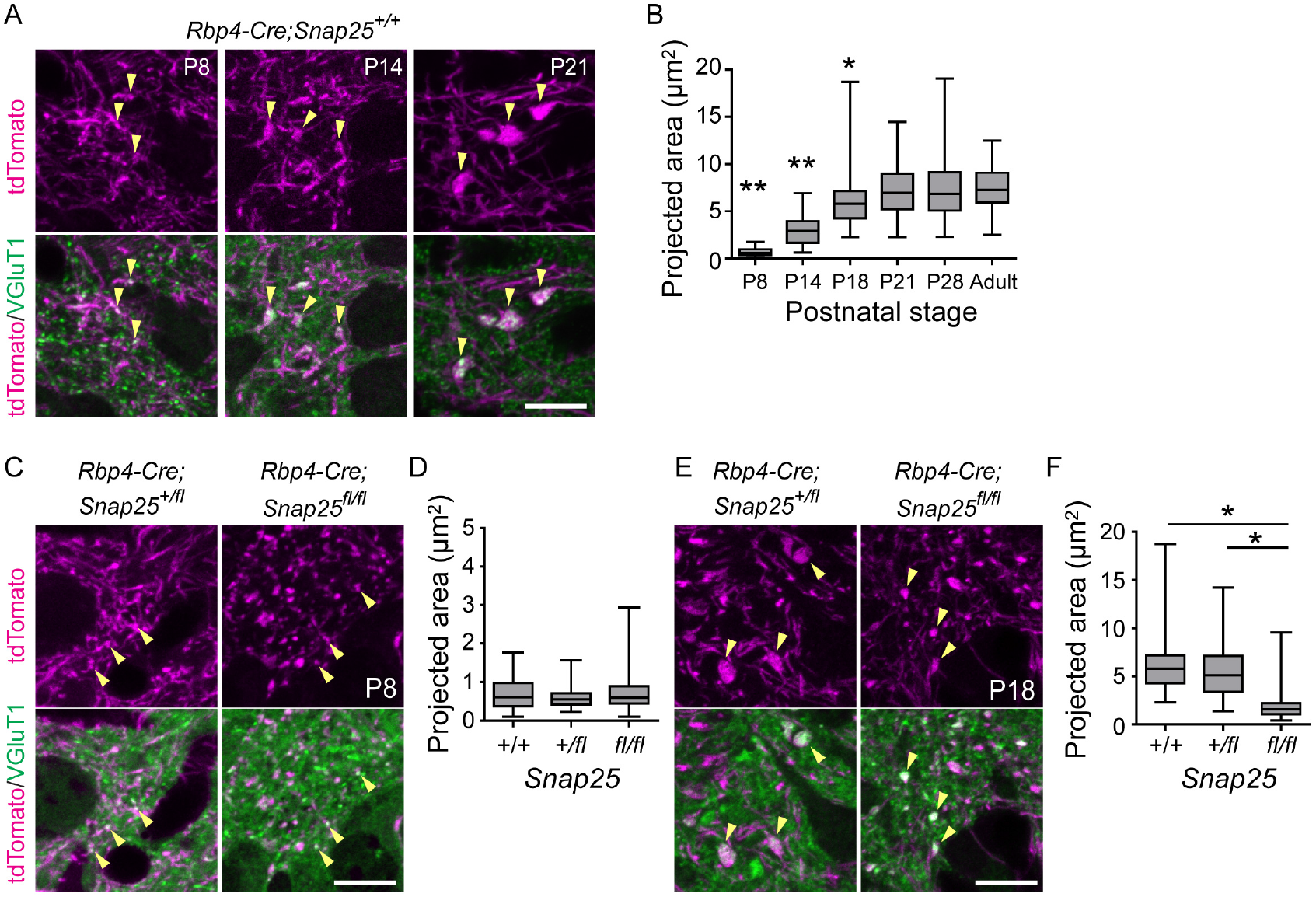
Development of *Rbp4-Cre+* layer 5 boutons in Po during the first four postnatal weeks. **A**, Laser-scanning confocal microscopy images of *Rbp4-Cre+* layer 5 boutons in Po of *Rbp4-Cre;Ai14* mice at P8 (left), P14 (middle) and P21 (right). Maximum intensity projections of z-stack images (1.0 μm for P8 and P14, 2.5 μm for P21 in total thickness) are shown. Arrows indicate representative *Rbp4-Cre* driven tdTomato+ boutons that are colocalised with VGluT1. **B**, Quantification of the size of tdTom+ boutons in projected images of z-stacks in mice during postnatal development. The size of boutons at P8-P28 were compared with those of adults. 90 boutons from 3 brains were measured for each stage. *, *P*<0.05; **, *P*<0.001. **C**, tdTom+ boutons in *Rbp4-Cre;Snap25^+/fl^* and *Rbp4-Cre;Snap25^fl/fl^* at P8. **D**, There were no statistically significant differences between *Snap25^+/+^*, *Snap25^+/f^* and *Snap25^fl/fl^* genotypes when comparing projected area size of VGluT1+tdTom+ boutons. **E**, tdTom+ boutons in *Rbp4-Cre;Snap25^+/fl^* and *Rbp4-Cre;Snap25^fl/fl^* at P18. **F**, the size of *Rbp4-Cre;Snap25^fl/fl^* boutons was significantly smaller than those of controls. n=90 boutons from three brains for each genotype were analysed in **D**,**F**. *, *P*<0.001. One-way ANOVA followed by the post hoc test with Dunn’s Multiple Comparison Test. Scale bars, 10 μm.

### SNAP25 is required for the normal development of layer 5 boutons in Po

We next sought factors that regulate the development of layer 5 giant boutons in Po. We focused on SNAP25, which is essential for regulated vesicular release at synapses (Washbourne et al., 2002). We had previously validated the lack of regulated synaptic vesicle release in the floxed *Snap25* mouse model in the presence of Cre recombinase expression *in vitro* and *in vivo* ((Hoerder-Suabedissen et al., 2019; Marques-Smith et al., 2016). This showed that there is no difference in the initial projections of *Rbp4-Cre*; *Snap25^fl/fl^* neurons into Po (Hoerder-Suabedissen et al., 2019). We found no difference in the size of the tdTom+ boutons at P8 between *Rbp4-Cre*; *Snap25^fl/fl^* and controls (*Rbp4-Cre*: *Snap25^+/+^* and *Rbp4-Cre*; *Snap25^+/fl^*) (Fig. 2C, D). At P18, however, the size of boutons in *Rbp4-Cre*; *Snap25^fl/fl^* was significantly smaller than that of controls (Fig. 2E, F). To understand what this size difference means in terms of their structure we performed ultrastructural analysis of knockout boutons using post-embedding immuno-electron microscopy to identify those with tdTomato. At P8, tdTom+ boutons formed a synapse with Po dendrites in randomly selected EM sections of both control and *Rbp4-Cre*; *Snap25^fl/fl^* mice (Fig. 3A). At this age, there were no significant differences in the bouton size or the number of synapses per bouton in control and *Rbp4-Cre*; *Snap25^fl/fl^* mice (Fig. 3B,C). By P18, however, excrescences of Po dendrites showed multiple synapses with tdTom+ boutons in *Rbp4-Cre*; *Snap25^+/+^* brains (Fig. 3D left). In contrast, *Rbp4-Cre*; *Snap25^fl/fl^* boutons did not envelop excrescences and only showed a single synapse (Fig. 3D right). The size of boutons, the area of excrescences, and the number of synapses per bouton, per section, were significantly smaller in *Rbp4-Cre*; *Snap25^fl/fl^* compared to controls (Fig. 3E-G). These results indicate that SNAP25 in presynaptic neurons is required for the maturation of the excrescences on the dendrites of Po neurons, and the subsequent formation of multiple synapses.

**Figure 3.**
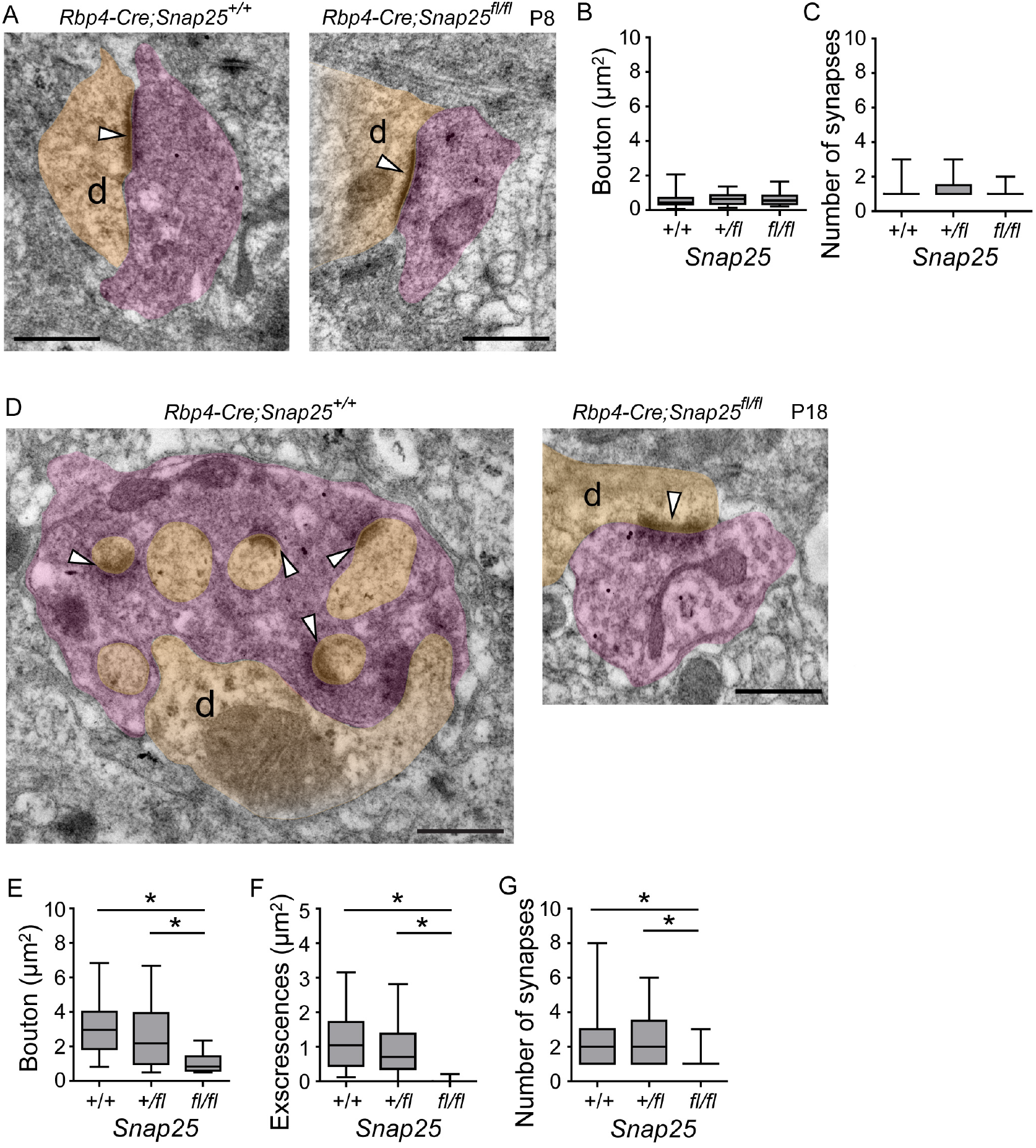
*Rbp4-Cre;Snap25^fl/fl^* boutons lack excrescences from Po dendrites at P18. **A-C**, Ultrastructure obtained by post-immunoelectron microscopy of *Rbp4-Cre;Snap25^+/+^* and *Rbp4-Cre;Snap25^fl/fl^* synapses at P8, which do not contain any excrescences from the associated postsynaptic compartment (**A**). There are no statistical differences between the two in the size of boutons or the number of synapses (**B**,**C**). **D-G**, Ultrastructure of *Rbp4-Cre;Snap25^+/+^* and *Rbp4-Cre;SNAP25^fl/fl^* at P18, synapses with clear postsynaptic densities are indicated with white arrowheads (**D**). The size of boutons, the area of excrescences and the number of synapses are significantly decreased in *Rbp4-Cre;Snap25^fl/fl^* compared to controls (**E**-**G**). Purple and light orange indicate presynaptic boutons and postsynaptic dendrites, respectively, ‘d’ denotes the dendrite. One-way ANOVA followed by the post hoc test with Dunn’s Multiple Comparison Test, *, *P*<0.001; Scale bars, 500 nm.

### Presynaptic SNAP25 is required for the development of the glomerular type junction between layer 5 boutons and Po dendrites

To better understand the effect that SNAP25 removal has on the structure of these junctions between layer 5 boutons and Po dendrites we used correlative light and block face scanning electron microscopy (SBEM); reconstructing in 3D the boutons and excrescences at P18. Each *Rbp4-Cre*; *Snap25^+/+^* bouton contained multiple excrescences to which it was synapsing, all of which were derived from a single dendrite. The bouton completely enveloped the excrescences, and glial processes covered the boutons (Fig. 4A,B). Reconstruction of individual excrescences showed their diverse morphology such as straight or curved shape with or without branching (Fig. 4C). Each process, or spine, of the excrescence are packed into a single bouton, each being independently isolated within the bouton, and unopposed to any other process of the same excrescence. In contrast, *Rbp4-Cre*; *Snap25^fl/fl^* boutons did not contain dendritic excrescences and had a single synapse with the shaft of the connecting Po dendrite (Fig. 4D,E). *Rbp4-Cre*; *Snap25^+/+^* boutons contained 6.1±2.8 (mean±s.d. n=11 boutons from one brain) excrescences, whereas *Rbp4-Cre*; *Snap25^fl/fl^* boutons contained no excrescences (mean±s.d., n=11 boutons from one brain, Table). The surface of boutons in contact with the dendrite was greatly reduced in *Rbp4-Cre*; *Snap25^fl/fl^* (1.1±1.4 μm^2^, mean±s.d., Table) compared with that in *Rbp4-Cre*; *Snap25^+/+^* (41.5±12.7 μm^2^ mean±s.d., Table). The total and net volume of boutons with and without including excrescences, in *Rbp4-Cre*; *Snap25^+/+^* were 9.2 ± 2.3 μm^3^ and 5.7 ± 1.5μm^3^, respectively (mean±s.d., n=11 boutons from one brain, Table). Both of those volumes are larger than the volumes in *Rbp4-Cre*; *Snap25^fl/fl^* (2.2 ± 1.4 μm^3^ mean±s.d., n=11 boutons from one brain, Table). Those results suggest that loss of excrescences in the *Rbp4-Cre*; *Snap25^fl/fl^* boutons results in reduction of the contact surface area and the number of synapses between the boutons and Po dendrites.

**Figure 4.**
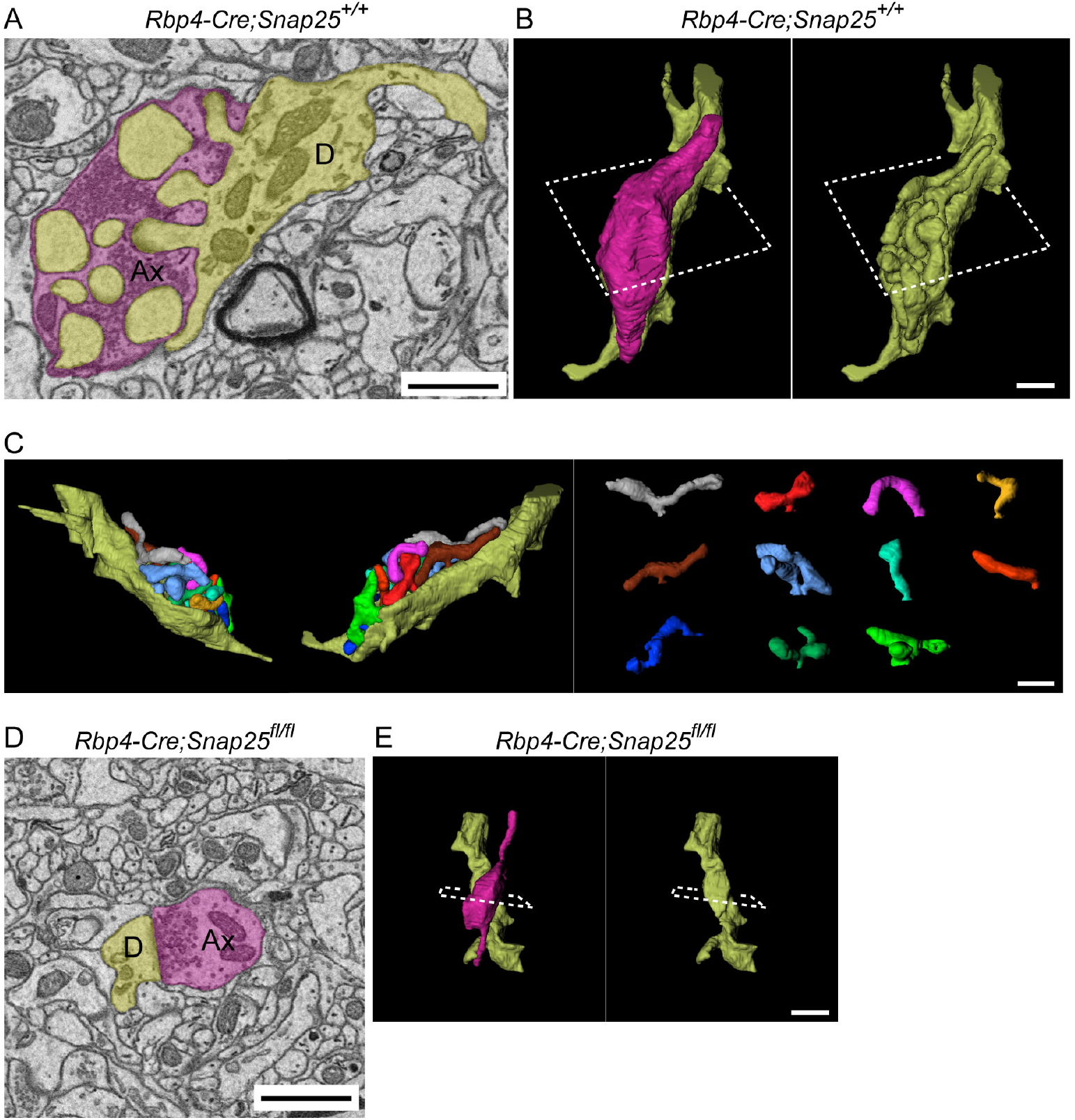
SBEM shows the structure of *Rbp4-Cre;Snap25^+/+^* and *Rbp4-Cre;Snap25^fl/fl^* boutons and their connected dendrites at P18. **A,B**, A single image from the series taken with the SBEM (**A**) and reconstructed 3D model (**B**) from Po in a *Rbp4-Cre;Snap25^+/+^* brain (Ax, axon; D, dendrite). **C**, Reconstruction of individual excrescences on the same dendrite shown in **A**,**B**. Each excrescence is shown with different colour in the right panel. **D**,**E**, A typical SBEM image (**D**) and reconstructed 3D model (**E**) from Po in an *Rbp4-Cre;Snap25^fl/fl^* brain. Scale bars, 1 μm.

### SNAP25 is required for the formation of normal synaptic structures

We compared the ultrastructure of synapses in Po between *Rbp4-Cre*; *Snap25^+/+^* and *Rbp4-Cre*; *Snap25^fl/fl^* at P18 by pre-embedding immuno-electron microscopy (Fig. 5A-G). In *Rbp4-Cre*; *Snap25^+/+^* brains, synaptic vesicles in tdTom+ giant boutons were round without dense-core (Fig. 5A, A’). The diameters of those vesicles were significantly larger than those of asymmetrical synapses formed by tdTom-negative (tdTom-) small boutons, which were found near the tdTom+ giant boutons analysed (Fig. 5B, E, left two plots, n=53 and 41 vesicles in tdTom+ and tdTom-presynapses, respectively from one brain). In contrast, the thickness of postsynaptic densities in tdTom+ synapses were significantly smaller than those in tdTom-synapses (Fig. 5F, left two plots, n=17 tdTom+ and 13 tdTom-synapses from one brain). The distance of their synaptic clefts showed no significant difference between the two (Fig. 5G, left two plots). These results suggest that tdTom+ giant boutons have a different synaptic profile from tdTom-small boutons in the size of vesicles and postsynaptic densities. In *Rbp4-Cre*; *Snap25^fl/fl^* brains, tdTom+ (*Snap25* cKO) presynapses also contained round vesicles without dense-core (Fig. 5C, C’) and those vesicles were significantly larger than those of asymmetrical synapses formed by tdTom-small boutons (Fig. 5D, E, right two plots, n=67 and 42 vesicles in tdTom+ and tdTom-presynapses, respectively from one brain). Those vesicles in tdTom+ presynapses in *Rbp4-Cre*; *Snap25^fl/fl^* were even larger than the counterparts in *Rbp4-Cre*; *Snap25^+/+^* (Fig. 5E, 1st and 3rd plots from the left). Postsynaptic densities of tdTom+ synapses in *Rbp4-Cre*; *Snap25^fl/fl^* were also significantly larger than the counterparts in *Rbp4-Cre*; *Snap25^+/+^* (Fig. 5F, 1st and 3rd plots from left, n=17 and 23 synapses, respectively in *Rbp4-Cre*; *Snap25^+/+^* and *Rbp4-Cre*; *Snap25^fl/fl^*), whereas there was no difference between the two in the distance of their synaptic clefts (Fig. 5G, 1st and 3rd plots from the left). These results show that as well as changing the bouton morphology, deletion of pre-synaptic SNAP25 also alters some of the synaptic profiles of the *Rbp4-Cre*+ layer 5 giant boutons.

**Figure 5.**
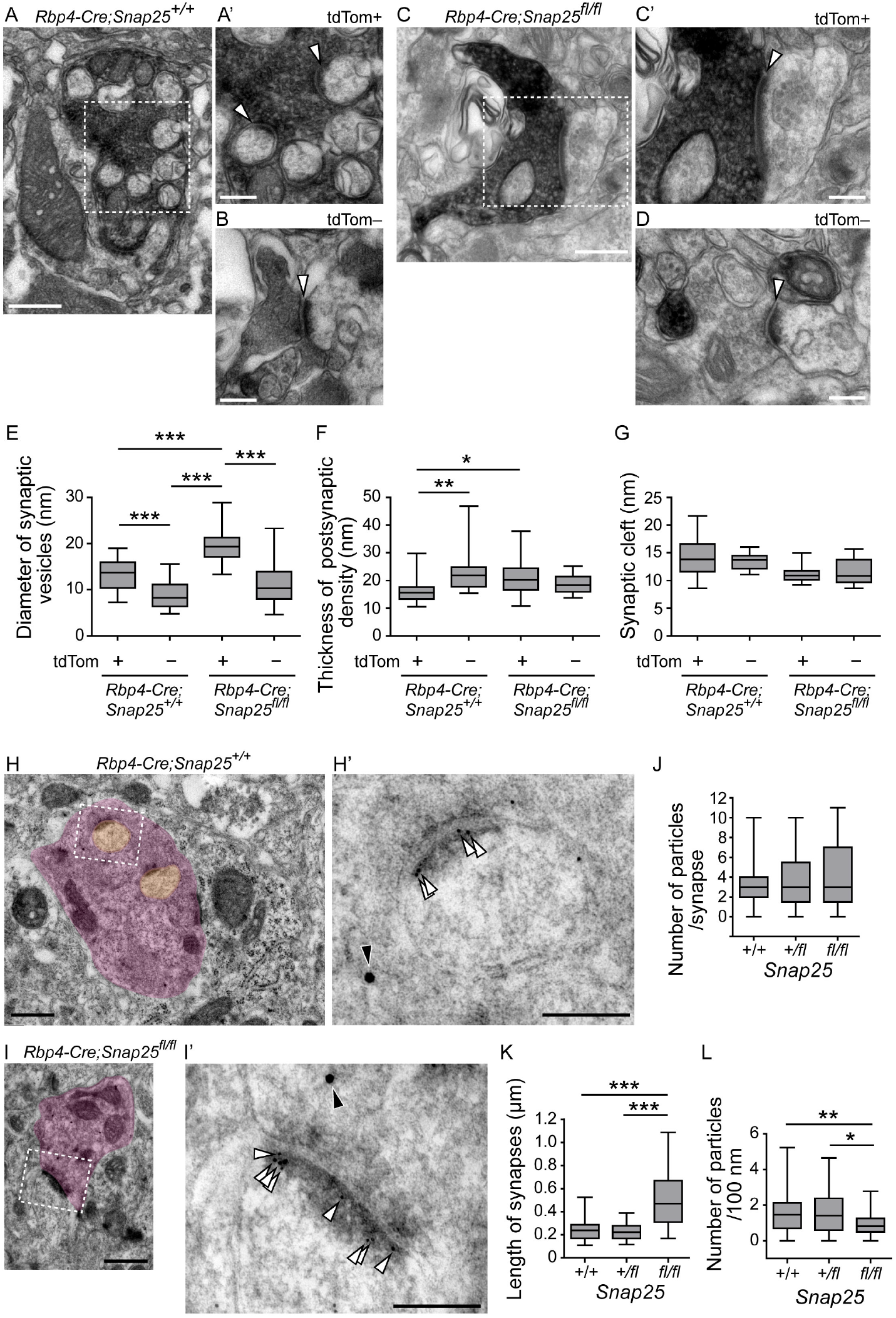
Assembly of synapses between *Rbp4-Cre+* layer 5 boutons and Po dendrites in control and *Rbp4-Cre;Snap25^fl/fl^* brains at P18. **A**, **B**, Electron micrographs obtained by pre-embedding immuno-electron microscopy against tdTomato showing synapses formed by *Rbp4-Cre*-driven tdTomato-positive (tdTom+) (**A**,**A**’) or tdTom-negative (tdTom-) boutons (**B**) with Po dendrites in *Rbp4-Cre;Snap25^+/+^* brains. **A’** is a magnified image of the boxed region in **A**. Arrowheads indicate synapses. **C**, **D**, Electron micrographs showing synapses formed by tdTom+ (**C**,**C**’) or tdTom-boutons (**D**) with Po dendrites in *Rbp4-Cre;Snap25^fl/fl^* brains. **C’** is a magnified image of the boxed region in **C**. Arrowheads indicate synapses. **E**-**G,** The diameter of synaptic vesicles in presynapses (**E**), the thickness of postsynaptic density (**F**) and the distance of synaptic cleft (**G**) of tdTom+ and tdTom-synapses in *Rbp4-Cre;Snap25^+/+^* and *Rbp4-Cre;Snap25^fl/fl^* brains. **H**, **I**, Ultrastructure of an *Rbp4-Cre+* layer 5 bouton and its connecting dendrite in *Rbp4-Cre;Snap25^+/fl^* (**H**) and *Rbp4-Cre;Snap25^fl/fl^* (**I**). 10 nm and 20 nm immuno-gold particles were used to detect PSD95 and tdTomato, respectively by post-embedding immunoelectron microscopy. **H’** and **I’** are a magnified image of the boxed region in **H** and **I**, respectively. White and black arrowheads indicate 10 nm and 20 nm immuno-gold localisation, at postsynaptic density **J**-**L**, Length of synapses (**J**), the number of immuno-gold particles per synapse (**K**) and the number of particles per 100 nm (**L**) were compared in the three genotypes. n=52, 50 and 34 synapses from three brains for each of *Rbp4-Cre;Snap25^+/+^, Rbp4-Cre;Snap25^+/fl^, Rbp4-Cre;Snap25^fl/fl^*. *, *P*<0.05; **, *P*<0.01; ***, *P*<0.001. Scale bars, 500 nm in **A**, **C**, **H** and **I**, 200 nm in **A’**, **B**, **C’, D, H’** and **I’**.

We also compared the localisation of PSD95 as a marker of postsynaptic densities by post-embedding immuno-electron microscopy of tdTom+ boutons in control animals (*Rbp4-Cre*; *Snap25^+/+^* and *Rbp4-Cre*; *Snap25^+/fl^*) and *Rbp4-Cre*; *Snap25^fl/fl^*. PSD95 localised at postsynaptic densities on excrescences of Po dendrites that synapse with tdTom+ positive boutons in control at P18 (Fig. 5H, H’). Postsynaptic densities on Po dendrites that synapse with tdTom+ (*Snap25* cKO) boutons in *Rbp4-Cre*; *Snap25^fl/fl^* brains also contained PSD95 (Fig. 5I, I’). The number of immuno-gold particles per synapse was not significantly different between control and *Rbp4-Cre*; *Snap25^fl/fl^* (Fig. 5J). However, the length of synapses was significantly larger in *Rbp4-Cre*; *Snap25^fl/fl^* than in control (Fig. 5K). This resulted in the decrease of the density of particles (the number of particles per 100 nm) in *Rbp4-Cre*; *Snap25^fl/fl^* compared with that in control (Fig. 5L). These results support the idea that assembly of synapse structure itself is not much affected, but some synaptic profiles are altered by SNAP25 removal from presynapses of *Rbp4-Cre*+ layer 5 giant boutons.

## Discussion

We show that the maturation of specialised junctions between cortical layer 5 axons from somatosensory cortex and Po thalamic dendrites requires the presence of SNAP25. Morphologically, the initial synapse formation of *Rbp4-Cre* positive layer 5 corticothalamic projections in Po does not appear to be affected by the absence of SNAP25. However, their subsequent maturation, with the formation of excrescences of Po dendrites, envelopment by the bouton to form a single glomerulus is suppressed. These results suggest that SNAP25-mediated regulated vesicular release from presynaptic terminals plays a crucial role in coordinated morphogenesis of the pre- and post-synaptic structure of layer 5 corticothalamic projections in Po.

### Ultrastructure of the interface between layer 5 boutons and Po dendrites

By using immuno-electron microscopy we have shown that the synaptic development of layer 5 corticothalamic projections in Po takes two steps: the axons form small boutons that synapse on the shaft of Po dendrites by the end of the first postnatal week, and then the boutons become larger in the second and third postnatal weeks. Each of those large boutons contains ~10 excrescences from Po dendrites. Mossy fibres from the hippocampal dentate gyrus also form characteristic large boutons that synapse with thorny spines on CA3 dendrites (Rollenhagen and Lubke, 2010). The mossy fibre boutons have 2 to 13 μm^3^ in volume and this size is comparable to that of layer 5 boutons in Po. Both boutons contain numerous mitochondria. However, there is also a difference between the two giant boutons: the mossy fibre giant boutons have filopodial extensions that form contacts with GABAergic interneurons and CA3 dendrites (Acsády et al., 1998; Martin et al., 2017), whereas no such extensions were observed in layer 5 giant boutons in Po. Compared to the mossy fibre boutons, the layer 5 boutons appear to wrap all the connecting excrescences more completely and glial processes further cover the bouton.

*Snap25* wild-type boutons have an area of contact with the connecting Po dendrites that is 35 times larger than its knockout counterpart. This shows a massive expansion of the contact surface up to the P18 time point with the formation of the excrescence structures. However, the electron microscopy indicated large synapses formed by *Rbp4-Cre*; *Snap25^fl/fl^* boutons, although the amount of gold-labelled PSD95 protein per synapse was not different from that of wild-type. This led to a reduced density of PSD95 signals in *Rbp4-Cre*; *Snap25^fl/fl^* synapses and raised the possibility that an expansion of the synapse is an adaptive change to deficient synaptic transmission (Goel et al., 2019; Murthy et al., 2001).

### Differential effects of presynaptic silencing on postsynaptic dendritic structures

Our results have shown that the initial synaptic assembly is not affected by *Rbp4-Cre*; *Snap25^fl/fl^*. This suggests that synaptic communication via SNAP25 is dispensable for the assembly of the initial synapses but essential for the subsequent establishment of specialised synaptic structure with the protrusion of dendritic excrescences and layer 5 bouton invaginations. Importantly, in the *Rbp4-Cre*; *Snap25^fl/fl^* brains we used, *Snap25* was removed only in presynapses of layer 5 giant boutons in Po, not the Po neurons themselves. Lack of excrescences on Po dendrites in those brains therefore suggests that presynaptic vesicular release is essential for the morphological changes of post-synaptic structures in Po neurons. Normal assembly of synapses under silencing is in agreement with previous observations obtained by other silencing methods in the hippocampus (Lu et al., 2013; Sando et al., 2017; Sigler et al., 2017). This suggests the initial assembly of synapses is regulated by release independent mechanisms such as synaptic organising cell adhesion molecules (Siddiqui and Craig, 2011), but the subsequent maturation and/or further morphological changes of synaptic contact sites such as formation of excrescences require evoked synaptic vesicle release or vesicle secretion from presynapses. However, the previous studies reported that there is a reduction in the number of synapses in the cortex of *Munc18-1* null mutants (Bouwman et al., 2004) and also that Emx1-driven TeNT causes a significant decrease of dendritic arborisation and ~ 40% decrease of spine densities in the CA1 (Sando et al., 2017). Moreover, a study using live imaging of hippocampal organotypic slice culture showed that blocking postsynaptic NMDA and AMPA receptors with antagonists reduces recruitment of PSD95 to newly formed spines and the stability of spines (De Roo et al., 2008). Decreased recruitment of PSD95 was also observed in retinal ganglion cells (RGCs) in the synapse formed by bipolar cells that express TeNT (Kerschensteiner et al., 2009). Those results suggest that synaptic transmission may be involved in stabilisation of newly formed spines, but the impact of silencing can be varied depending on the neuronal circuit analysed and the method of perturbing synaptic communication.

### Developmental regulation of giant bouton formation

Multiple synaptic contacts between single presynaptic boutons and dendrites are also found in other thalamic nuclei including dorsal lateral geniculate nucleus (dLGN), in which axons from retinal ganglion cells (RGCs) form large boutons (Bickford, 2015; Guillery, 1969; Morgan et al., 2016). In the dLGN, the bouton size is initially small at P7 and large boutons emerge by P14 (Bickford et al., 2010). This timing of giant bouton formation in dLGN is similar to that of layer 5 boutons in Po. The regulatory mechanism for the development of those large boutons is not fully understood. In hippocampal mossy fibres, postsynaptic ligand of Numb protein X 1 (Lnx1) and EphB receptors retrogradely regulate the fibre terminal maturation (Liu et al., 2018). It has also been shown that the interaction between the heparan sulfate proteoglycan GPC4 and the orphan receptor GPR158 organise mossy fibre-CA3 synapses (Condomitti et al., 2018). It would be of interest to determine whether synaptic vesicular release via SNAP25 regulates the molecular pathways involved in the mossy fibre bouton formation. However, it could also be possible that the mechanism underlying postsynaptic morphological changes is different between mossy fibre-CA3 and layer 5-Po synapses. A previous study reported that formation of thorny spines were 30% increased than control by Emx1-driven TeNT expression (Sando et al., 2017). In *Gpr158* knockout mice, thorny spine formation in mossy fibre-CA3 appears not to be greatly disturbed (Condomitti et al., 2018). Comparing the effect of silencing of presynaptic neurons with the same method between mossy fiber-CA3 and layer 5-Po systems would provide further insights into the underlying mechanism for the specialised synaptic structures in the two systems.

Our results clearly indicate that activity mediated through regulated vesicular release from the presynaptic terminal is not necessary for the formation of synapse, but it is required for the establishment of glomerular structures between layer 5 corticothalamic projections in Po. While the previous publications that reported no change in synapse formation used organotypic slice cultures of double knockout of *Munc13-1* and *13-2* (Sigler et al., 2017) or Emx1-driven tetanus toxin (TeNT) expressing brains (Sando et al., 2017), we used conditional *Snap25* KO in a specialised synapse in vivo. The lack of regulated vesicular release had an obvious effect, but it is not clear how SNAP25-depenent synaptic vesicular release regulates morphological changes of layer 5 boutons and excrescence formation in Po. One possibility is that neurotrophic factors such as BDNF released via SNARE-dependent exocytosis from presynapses (Shimojo et al., 2015) activate postsynaptic signalling pathways to form excrescences from Po dendrites. The specialised synapses between Layer 5 and Po dendrites present excellent model systems to further investigate the molecular mechanisms underlying SNAP25-dependent specialised synaptic development of layer 5 corticothalamic projections in Po.

## Materials and Methods

### Breeding and Maintenance of Transgenic Mice

The animal experiments were performed in the Biomedical Services of the University of Oxford (UK) under a Animals Scientific Procedures Act 1986 project licence as well as with local ethical approval by the central Committee on Animal Care and Ethical Review (ACER) and the Animal Welfare and Ethical Review Body (AWERB) at the University of Oxford. Tg(Rbp4-cre)KL100Gsat/Mmucd (Rbp4-Cre; Jackson Laboratories) mice were crossed with B6;129S6-Gt(ROSA)26Sortm14(CAG-tdTomato)Hze/J (Ai14) to label cortical layer 5 neurons. To generate *Rbp4-Cre*; *Snap25^fl/fl^* mice, the above strain was crossed with B6-Snap25tm3mcw (*Snap25^fl/fl^*) mice, which were obtained from University of New Mexico (Michael C. Wilson) (Hoerder-Suabedissen et al., 2019).

### Cre-dependent Adeno Associated (AAV) Viral injection to somatosensory cortex

To trace the axonal projections of Cre+ L5 axons in Po thalamus, AAV2-CAG-Flex-ArchT-GFP (University of North Carolina Vector core) was injected in primary somatosensory cortex (S1) of *Rbp4-Cre;Ai14* young adult animals (6 weeks) old. Mice were deeply anaesthetized with isoflurane, and placed in a stereotaxic frame. Following midline skin-incision, a craniotomy was performed over right hand side S1 cortex (1.5 caudal, 2.75-2.8 lateral). Virus-filled pulled glass-micropipettes were inserted into the brain to the required depth (0.6-0.7) and 200 nl of virus were slowly pressure ejected into the brain. Pipettes were retracted 5 min after the last ejection of virus, followed by post-surgery repair and recovery of the animal. Post-injection survival was ranged 3 weeks, at which point animals were terminally anaesthetized and perfusion-fixed with 4% paraformaldehyde (PFA, Sigma-Aldrich, F8775) and 0.2% glutaraldehyde (Electron Microscopy Sciences, 16220) in phosphate buffer (PB) at pH7.4 as described in Pre-embedding immunoelectron microscopy section below.

### Immunohistochemistry

Young animals at 1-4 weeks age and 3-4 month old adults were perfusion fixed with 4% formaldehyde in 0.1M PBS, and dissected brains were postfixed in the same solution for 24 hrs at 4 °C. Brains were sectioned coronally at 50 μm on a vibrating microtome (Leica, VT1000S). Sections were incubated in PBS containing 2% goat serum or 3% BSA (blocking solution) for 2 hrs, followed by incubation of anti-Vesicular Glutamate Transporter 1 (VGluT1) antibody (1:2000, Millipore, AB5905) in the blocking solution for overnight at 4°C. Staining was visualised by secondary antibodies conjugated with Alexa 488, Alexa 568, Cy3, Alexa 633 or Cy-5 (Thermo Fisher). Sections were counterstained with 4’,6-diamidino-2’-phenylindole dihydrochloride (DAPI) to visualise the nuclei.

### Serial Block Face Scanning Electron Microscopy (SBEM)

SBEM was performed as previously described (Maclachlan et al., 2018). Briefly, *Rbp4-Cre;Ai14;Snap25^+/+^* and *Rbp4-Cre;Ai14;Snap25^fl/fl^* brains at P18 were perfused with 0.1 M phosphate buffer containing 2% PFA (Electron Microscopy Sciences, 15714) and 2.5% glutaraldehyde (Electron Microscopy Sciences, 16220), at pH7.4, and post-fixed at room temperature for 2 hrs. Dissected brains were vibratome sliced at 80 μm and fluorescent and bright-field images of Po regions at various magnifications collected using an epifluorescence microscope (Leica DMR) and confocal microscope (LSM710 Zeiss). The position of tdTom+ boutons were imaged along with major blood vessels as fiducial marks so that boutons in fluorescence images could be localised in electron micrographs by their location with respect to these other features. The sections were postfixed in 1.5% potassium ferrocyanide (Sigma-Aldrich, 14459-95-1) and 2% osmium tetroxide mixed together (Electron Microscopy Sciences, 19110). They were then stained with 1% thiocarbohydrazide (Sigma-Aldrich, 101001342) followed by 2% osmium tetroxide and then stained overnight in 1% uranyl acetate (Electron Microscopy Sciences, 22400). The final stain was at 50 °C, in a lead aspartate solution at pH 5, washed in water, and infiltrated with Durcupan resin (Electron Microscopy Sciences). The sections were mounted between glass microscope slides coated in a mould separating agent (Glorex Inspirations, Switzerland, 6 2407 445) and the resin hardened at 65 °C for 24 hrs. The sample was imaged with a SEM microscope (Merlin, Zeiss NTS) fitted with the 3View cutting system (Gatan, Inc., Pleasanton, CA, USA). The obtained image series was aligned using the alignment functions in the TrakEM2 plugin of FIJI (Cardona et al., 2012) and segmented. The models were then exported into the Blender software and analysed using the NeuroMorph tools (Jorstad et al., 2018; Jorstad et al., 2015).

### Post-embedding immunoelectron microscopy

Mice were perfused with PB containing 2% PFA and 2.5% glutaraldehyde at pH7.4. Brains were coronally sectioned at 80 μm and the Po region was dissected out under a fluorescence microscope (MZFLIII, Leica). Tissue pieces containing Po were stained with 2% uranyl acetate (Agar Scientific, AGR1260A) in 0.1M sodium acetate buffer (Sigma-Aldrich, S2889) for 45 min–1 h and dehydrated through a graded series of methyl alcohol (70%, 90% and absolute) at −20 °C. The tissues were embedded in LR gold resin (Agar Scientific, AGR1284) containing 0.5% benzil (Agar Scientific, AGR1285) under UV light for 16–18 hrs at −20 °C. Ultrathin sections (70 nm) were prepared on an ultramicrotome (Leica Ultracut S) and mounted on 200-mesh nickel grids (Agar Scientific, AGG2200N) coated with formvar (TAAB, F145/025). For immunolabelling, sections were blocked with 1% chicken egg albumin (Sigma-Aldrich, A5503) in PBS and incubated with rabbit anti-red fluorescent protein (RFP) antibody (1:500, PM005, MBL international) for 2 h and then with 20 nm gold particle-conjugated goat anti-rabbit (1:50, BBI solutions, EM.GAR20) for 1 hr. For detection of PSD95, anti-PSD95 antibody (1:100, Synaptic Systems, 124 014) and 10 nm gold particle-conjugated goat anti-Guinea Pig (1:50, BBI solutions, EM.GAG10) were used. For negative control sections, the primary antibody was omitted. The sections were postfixed with 1% glutaraldehyde for 10 min and lightly counterstained with 2% uranyl acetate and 2.77% lead citrate solution (Agar Scientific, AGR1210). The immunolabelled sections were examined on a JEOL 1010 transmission electron microscope (JEOL) fitted with an Orius digital camera (Gatan).

### Pre-embedding immunoelectron microscopy

Mice were perfused with PB containing 4% PFA and 0.2% glutaraldehyde at pH7.4. The brains were cut at 50 μm-thick, and freeze and thaw of the slices were performed in 30% sucrose twice. For detection of AAV-eGFP, the slices were blocked in 1%BSA and incubated with anti-GFP antibody (1:500, Invitrogen, A-11122) for 24 h at 4°C followed by goat biotinylated anti-rabbit antibody (1:250, Vector Laboratories, BA-1000) for 2h at RT. Signals were enhanced with the ABC method according to the manufacturer’s instruction (Vector Laboratories, PK-6100) and visualised with 3,3′-Diaminobenzidine tetrahydrochloride (DAB, Sigma-Aldrich, D3939) for 5 min at room temperature. The slices were postfixed with 3% glutaraldehyde and then 1% osmium tetroxide in PB for 1h at RT and stained with 2% uranyl acetate in H_2_O for 45 min-1h at RT. The slices were dehydrated through a graded series of ethyl alcohol (30%, 50%, 70%, 80%, 90% and absolute) at 4°C and replaced with acetone followed by Spurr low viscosity embedding medium, whose formula is as follows: ERL4221 (Agar Scientific, AGR1047R, 4.1 g), DER 736 Diglycidylether of Polypropyleneglycol (Agar Scientific, AGR1072, 0.95 g), Nonenyl Succinic Anhydride (NSA) (Agar Scientific, R1055, 5.9 g) and Dimethylaminoethanol (S1) (Agar Scientific, R1067, 0.1ml). The slices were embedded between Aclar film (Agar Scientific, AGL4458) and the resin was hardened at 60°C for 24h. 70 nm-thick ultrathin-sections were prepared and imaged with the same ultramicrotome and electron microscopy as in post-embedding electron microscopy.

For detailed analysis of synaptic profiles, samples were prepared and examined as described previously (Kiyokage et al., 2017). Briefly, brains were perfused and sectioned as described above, and sections were stained with rabbit anti-RFP antibody (1:1000, a kind gift from T. Kaneko) and then biotinylated goat anti-rabbit antibody (1:200, Vector Laboratories, BA-1000). Signals were enhanced with the ABC method followed by incubation in 0.05% DAB and 0.01% H_2_O_2_. After dehydration, the sections were flat-embedded in Epon-Araldite between a glass slide and a coverslip, both of which were pre-coated with a liquid-releasing agent (Electron Microscopy Sciences, 70880). Selected areas for EM observation were cut into 70-nm-thick serial sections with an ultramicrotome (Reichert-Nissei Ultra-Cuts, Leica) and examined with a transmission electron microscope (JEM-1400, JEOL).

### Image analysis and Statistics

Image J (https://imagej.nih.gov/ij/) software was used to analyse fluorescent and electron microscopy images. For the quantification of boutons in images obtained with laser scanning confocal microscopy, maximum intensity projection was applied to the z-stack fluorescent images (5 μm in total thickness) containing the whole volume of the bouton of interest, and the size of the bouton in the projected images was measured. For the quantification of the bouton in electron micrographs, boutons in randomly selected sections were analysed. Obtained data were analysed by one-way ANOVA followed by the post hoc test with Dunn’s Multiple Comparison Test using Prism 4 software (GraphPad). In the box-and whiskers graph, horizontal lines above, inside and below the box indicate maximum, median and minimum values, respectively and the top and bottom of the box indicate 75th and 25th percentiles, respectively.

## Author Contributions

SH and ZM conceived experiments. SH, AHS, GK and ZM wrote the manuscript. SH performed histological and ultrastructural analysis of boutons. AHS performed viral injections and helped supervise the project. CM and GK carried out the SBEM sample preparation and imagining. EK performed pre-embedding immunoelectron microscopy for analysis of synapses with advice from KT.

## Acknowledgements

We thank T. Kaneko for the anti-RFP antibody. ZM’s laboratory was supported by Medical Research Council (G00900901). SH was supported from Daiichi Sankyo Foundation of Life Science and The Uehara Memorial Foundation.

---

**Table.** Comparison of tdTom+ boutons and their connecting Po dendrites between *Rbp4-Cre*;*Snap25^+/+^* and *Rbp4-Cre*; *Snap25^fl/fl^* in Po at P18.

| | 1. Area of contact with<br>dendrite per bouton ($\mu\text{m}^2$) | 2. Number of<br>excrescences per bouton | 3. Volume of excrescences<br>per bouton ($\mu\text{m}^3$) | 4. Volume of<br>bouton ($\mu\text{m}^3$) | 5. Net volume of<br>bouton ($\mu\text{m}^3$)<sup>*1</sup> | 6. Number of<br>boutons analysed |
| --- | --- | --- | --- | --- | --- | --- |
| *Rbp4-Cre;*<br>*Snap25<sup>+/+</sup>* | 41.5 ± 12.7 | 6.1 ± 2.8 | 3.5 ± 0.9 | 9.2 ± 2.3 | 5.7 ± 1.5 | 11 |
| *Rbp4-Cre;*<br>*Snap25<sup>fl/fl</sup>* | 1.1±1.4 | 0 | 0 | 2.2 ± 1.4 | 2.2 ± 1.4 | 11 |

<sup>\*1</sup>, (5) = (4) – (3)

